# IMPROVED SSRs-BASED GENETIC DIVERSITY ASSESSMENT OF COCONUTS (*COCOS NUCIFERA* L) ALONG THE COAST OF KENYA

**DOI:** 10.1101/2021.10.01.462814

**Authors:** Justus C. Masha, Najya Muhammed, Vincent Njung’e, Maurice E. Oyoo, Manfred Miheso

**Affiliations:** Pwani University, P.O. BOX 195-80108, Kilifi, Kenya; International Crops Research Institute for the Semi-Arid Tropics (ICRISAT), P.O.BOX 39063-00100, Nairobi; Department of crops, Horticulture and soil sciences, Egerton University, P.O. BOX 536-20115, Egerton, Kenya; Kenya Agricultural and Livestock Research Organization - Food Crops Research Institute, Njoro, Kenya

**Keywords:** Coconut genetic diversity, capillary electrophoresis, polymorphic information content

## Abstract

Coconut is the most important cash crop along the Coast of Kenya, yet its genetic diversity has not been fully established. A genetic diversity study of 48 coconut genotypes that were collected along the Coast of Kenya was conducted with 13 polymorphic short sequence repeats (SSRs) markers. SSR analysis was performed using GeneMapper while data analysis was done with PowerMarker and DARwin softwares.

The results revealed a total of 68 alleles ranging from 2 to 11 per locus with a mean of 5.23 per marker. Gene diversity and polymorphic information content (PIC) ranged between 0.41 to 0.83 and 0.33 to 0.79, respectively. A neighbour-joining dendrogram grouped the genotypes into three major clusters containing distinct sub-clusters. This study underscored that capillary electrophoresis is a more accurate and informative technique for SSRs allele scoring than agarose gels, which was reported in a previous study with the same SSRs markers and coconut genotypes in Kenya. The clusters observed forms the basis to isolate conservation blocks, which are the key to establishing a genebank, since there is no documented coconut genebank for *ex-situ* conservation in Kenya.

## INTRODUCTION

Coconut (*Cocos nucifera* L.) is the main tropical cash crop and it provides a source of income to the people of the coastal lowlands of Kenya (Wekesa et al., 2017). Coconut is proposed to have originated from south Asia and disseminated initially to the Pacific and East African shores by floating on sea waves and later to the Atlantic and American shores by humans after cultivation (Harries, 1978). Consequently, in most studies, African and South Pacific coconut germplasm are closely related to those in the South Asian subcontinent (Lebrun et al., 1998; Teulat et al., 2000; Perera et al., 2003). The tall (up to 60 feet) and dwarf (up to 25 feet) coconut types as well as their intermediates, which are thought to be their hybrids (16 feet), are found at the coast of Kenya (Oyoo et al., 2015). Diversity is higher in Tall coconut varieties, which are outcrossing, than in the in-breeding Dwarf varieties (Teulat et al., 2000; Meerow et al., 2003). These varieties contribute significantly towards their social, economic and environmental wellbeing of the coastal people in the following counties where they are grown; Kwale, Kilifi, Mombasa, Tana River, Lamu and Taita Taveta. The coconut industry plays an important role in the protection of fragile environments such as small islands and coastal zones as well as providing a good tropical canopy for the tourism industry (Bourdeix and Prades., 2018).

Despite these diverse ecological services, the genetic diversity of coconut germplasm is under threat from climate change, pests and diseases, poor conservation strategies and urbanization, logging for timber and land fragmentation for housing, especially in coconut growing areas (Batugal et al., 2009; Martinez et al., 2009).

Assessment of the genetic diversity is an essential component of coconut genetic resource characterisation, genetic improvement and adoption of conservation strategies (George and Angels, 2008). For coconut, germplasm from different geographical areas might look similar but may be genetically different. They might also show morphological differences but may be genetically the same. Molecular marker technologies reduce the inclusion of duplicates in breeding programmes and conservation blocks (Batugal et al., 2009). Once the genetic diversity has been assessed using reliable DNA markers such as short sequence repeats (SSRs), distinct coconut representatives can be identified, collected and conserved in genebanks. Coconut diversity hotspots can be documented with precision and the richness of the genepool can be determined and regeneration of conserved accessions can be enhanced (Rao, 2004; Yao et al., 2013).

Molecular marker technology is an ideal tool for assessing coconut genetic diversity within and between coconut populations (Batugal et al., 2009). Genomic markers such as Random Amplified Polymorphic DNA (RAPD) (Masumbuko et al., 2014), Restriction Fragment Length Polymorphism (RFLP) (Lebrun et al., 1998), Amplified Fragment Length Polymorphism (AFLP) (Perera et al., 2000 and Teulat et al., 2000), Short Sequence Repeats (SSRs) or microsatellites (Dasanayaka et al., 2009; Martinez et al., 2009; Xiao et al., 2013) and inter-simple sequence repeats (ISSRs) (Manimekalai and Nagarajan, 2006) have been used to characterise coconut populations. Diversity in worldwide coconut germplasm show grouping patterns primarily according to dissemination routes from their source of origin (Rivera et al., 1999; Perera et al., 2003). SSR has been shown to be the most efficient and valuable molecular marker technology for genetic diversity studies and variety identification as are rich in polymorphism, high stability, and good repeatability (Dong et al., 2017)

Effective coconut genebanking strategies rely on analysis of genetic diversity at the DNA level in two ways: (1) to ensure that distinct coconut varieties from different geographical areas are represented in genebanks and (2) the DNA profiles serve as reference in regeneration of old coconut accessions for the maintenance of important agronomic traits [(Rao and Tobby (2004), Dasanayaka et al., 2009) and (Martial et al., 2013)].

The binary scoring method based on the presence or absence of bands (Wang et al., 2009), was deployed in the previous study (Oyoo et al., 2016). The genetic diversity of the same set of coconut germplasm using 13 SSRs markers failed to provide accurate genetic distances of the germplasm. Presently, the famous method for gene detection and isolation is polyacrylamide gel electrophoresis (PAGE) or agarose gel electrophoresis. However, these cannot reveal the accurate size of amplified target DNA fragments and has low detection efficiency. Furthermore, it is difficult to effectively integrate and accurately compare DNA fingerprinting data from large-scale samples and different batches of samples. Capillary electrophoresis based on DNA band and accurately determines the size of detected fragments thereby identifying subtle length differences. This study reported here employed the more informative and accurate capillary electrophoresis approach to resolve the multiple alleles that could be generated for a single marker across the 48 coconut genotypes and precisely sized them. In SSRs analyses, a missing amplification band does not indicate an absent SSRs allele. Furthermore, a visible band that appears in several different individuals are often several alleles with slightly different sizes (Mueller and Wolfenbarger, 1999). Some samples may present two different alleles for a single marker, indicating a heterozygous locus where the two diploid chromosomes each carry a different allele. Therefore, when SSRs markers are scored as presence or absence of alleles, such as when using agarose gels, their co-dominance and multi-allelic features are not considered, which can lead to misinterpretations (Jones et al., 1997; Wang et al., 2003).

The objective of this study was to describe the diversity of 48 coconut palms growing along the coast of Kenya using improved SSRs markers resolved by capillary electrophoresis.

## Materials and methods

### Area of study

The study is confined in the coastal region of Kenya which covers an area of approximately 82,383 km^2^ with a population of 4,329,474 (KNBS, 2019). The coastal region is comprised of 6 counties namely: Lamu, Kwale, Kilifi, Tana River, Mombasa and Taita Taveta. These counties in Kenya are known for their coconut diversity as reported in the previous study; Oyoo et al., (2015). The climate of this area is tropical humid with an annual mean rainfall of about 1200 mm mainly confined to the long rains between April to July and short rains between October and December (Mwachiro and Gakure, 2011).

### Sampling

Coconut leaf samples were obtained from the coastal region of Kenya in the following four counties; Kwale (GPS), Kilifi (GPS), Tana River (GPS) and Lamu (GPS). A total of 48 individual coconut genotypes previously collected by Oyoo et al., (2015) comprising of; 37 Tall, 8 Semi-tall (hybrids) and 3 Dwarf types. They were from different agro-ecological zones of the Coastal Lowlands of Kenya. Leaf samples were collected and coded according to the county of collection, district, division, sub-division, collection number and village. Sample collection was designed to account for differences in palm morphology, altitude, cropping systems and the local names.

### DNA Extraction and Evaluation

DNA was extracted from dry, frozen coconut leaves (preserved in silica gel at −20 °C) of the 48 varieties (tall, dwarf and hybrids) using a modified CTAB protocol described by Doyle and Doyle (1987). Two steel balls were placed in 2 mL labelled eppendorf tubes for each sample. Approximately 80 mg of dry leaves were weighed from each sample, cut into small pieces and ground into a fine powder to increase surface area for detergent activity, using a Tissue Lyser II (Qiagen^®^). This was followed by incubation with 800 μL of preheated CTAB extraction buffer (3 % CTAB w/v, 100 mM Tris-HCI (pH 8.0), 1.4 M NaCl, 20 mM EDTA, 3 % β-mercaptoethanol (BME), 2 % Polyvinylpyrollidone (PVP)(w/v) in a water bath at 65 °C for 30 minutes with occasional mixing. BME 3 % was used in a replica experiment to determine if it reduced the degree of degradation. Solvent extraction was done by adding 800 μl of chloroform:isoamyl alcohol (24:1) followed by thorough mixing. They were then centrifuged for 10 minutes at 13000 rpm and approximately 500 μL of the supernatant was transferred into clean labelled tubes. The DNA was precipitated by addition of 350 μL of isopropanol (0.7 volume) stored at −20 °C, left overnight to increase precipitation and then centrifuged for 20 minutes. The supernatant was decanted and the DNA pellet washed with 400 μL of 70 % ethanol, air dried for 30 minutes and re-suspended in 200 μL of low salt TE buffer (10 mM Tris-HCl pH 8.0, 1 mM EDTA). RNase A (5 μL of 10 mg/mL) was added and the samples were incubated at 37 °C for 1 hour to digest all RNA. The DNA was precipitated by the addition of 315 μL of ethanol sodium acetate (Ethanol: 3 M NaOAc 300 μL:15 μL) and incubated for 2 hours at −20 °C. The samples were centrifuged at 14000 rpm for 25 minutes, the supernatant was decanted and the pellet was washed with 400 μL of 70 % ethanol at 14000 rpm for 5 minutes. The DNA was air dried for 1 hour in a laminar flowhood, re-suspended in 50 μL low salt 1×TE and stored at −20 °C

### Evaluation of quality and quantity of the genomic DNA

The quality of DNA was evaluated by electrophoresis using 0.8 % (w/v) agarose gels stained with 5 μL/100 ml Gel Red^®^ (Biotium Inc., USA) to enable visualization. A mixture of 4 μl of DNA and 2 μl of loading buffer (25 mg bromophenol blue (0.25 %), 25 mg xylene xyanol (0.25 %), 4 g sucrose (40 %), was loaded onto the gel and run for 1 hour at 80 volts in a 0.5 × TBE buffer (0.1 M Tris base, 0.1 M boric acid and 0.02 M EDTA; pH 8.0). The fragments were visualized under UV light and photographed using a transilluminator. The amount and purity of the DNA quantity was determined by spectrophotometry using a Nanodrop^©^ 1000 (Thermo Scientific, USA). Programmed to measure absorbance (A) from; 220 to 350 nm and display the DNA concentration according to Wilfinger et al., (1997).

### PCR Amplification

PCR reactions were conducted in 10 μl final volume containing 1 x PCR buffer (20 mM Tris-HCl (pH 7.6); 100 mM KCl; 0.1 mM EDTA; 1 mM DTT; 0.5 % (v/v) Triton X - 100; 50 % (v/v) glycerol), 2 mM MgCl2, 0.16 mM dNTPs, 0.16 μM of a labelled M13-primer, 0.04 μM M13 - forward primer (InqabaBiotec, South Africa), 0.2 μM reverse primer (InqabaBiotec^™^), 0.2 units of Taq DNA polymerase (SibEnzyme Ltd, Russia), and 20 ng of template DNA as shown in Table 1

**Table 1:** Summary of concentration and volumes for individual PCR reagents per reaction

| Master mix components | Stock | Final concentration | Volume (µL) for 1 reaction |
| --- | --- | --- | --- |
| PCR Buffer without MgCl<sub>2</sub> | 10 x | 1 x | 1.0 |
| MgCl<sub>2</sub> | 50 Mm | 2 Mm | 0.4 |
| DNTPs | 2Mm | 0.16 Mm | 0.8 |
| M13 fluorescent Forward prime | 2 µM | 0.16 µM | 0.8 |
| Forward primer | 2 µM | 0.04 µM | 0.2 |
| Reverse primer | 2 µM | 0.2 µM | 1.0 |
| Taq polymerase | 5 U | 0.2 U | 0.2 |
| Template DNA |  | 20 ng/ µL | 1.0 |
| Sterile water |  |  | 4.6 |
| **Total volume** |  |  | **10** |

A total of 30 pairs of SSRs primers (supplementary Table 2) developed by Perera et al. (2000) were used in this study. For detection of PCR fragments during capillary electrophoresis, each forward primer was labeled with one of the three 6-Carboxyfluorescein fluorescent dyes: 6-FAM^®^, 6-VIC ^®^ or 6-PET^®^ (Life Technologies Corporation, Carlsbad, USA). During capillary electrophoresis, the amplification products passed through a detection window and a light excited the fluorescent dye. The fluorescence was then visualized using a computer programme as relative fluorescent units (RFU) against fragment length in base pairs. An allele was scored for each data point as length in base pairs at the highest RFU peak.

Reactions were performed on a thermocycler (GeneAmp PCR system 9700^®^, Applied Biosystems, USA). Initially, the thermocycler was progammed with an initial denaturation of 94 °C for 5 minutes, followed by 35 cycles of 95 °C for 30 seconds, 57 °C for 1 minute, and elongation at 72 °C for 2 minutes. Final elongation was done at 72 °C for 20 minutes, and PCR products held at 15 °C. Since the markers had a range of annealing temperatures from 56 - 46 °C, this protocol did not work well for all primer sets and a gradient PCR protocol was introduced. The thermocycler was programmed with an initial denaturation of 94 °C for 3 minutes followed by 16 touch down cycles of 62 °C annealing for 15 seconds and 72 °C for 15 seconds in every cycle, with the annealing temperature decreased by 1 °C in each subsequent cycle, generating a range of annealing temperatures from 62 °C to 46 °C. This was followed by 21 cycles of 95 °C for 15 seconds, 58 °C for 15 seconds and 72 °C for 30 seconds. Final chain elongation was done at 72 °C for 7 minutes and reaction products held at 15 °C. To amplify markers that did not succeed with this protocol, the touch down cycles were increased from 10 to 15 cycles and the annealing time increased by 15 seconds. The protocol is summarised in Table 2 and was adopted after optimization of the annealing temperatures.

**Table 2:** Touch down PCR conditions.

| PCR touchdown programme |  |  |  |
| --- | --- | --- | --- |
|  | Step | Temperature/ °C | Time/sec |
| 16 Cycles | 1st denaturation | 94 | 180 |
|  | 2nd denaturation | 94 | 30 |
|  | Annealing | 62 – 1 for each cycle | 15 |
|  | Elongation | 72 | 15 |
| 25 Cycles | Denaturation | 95 | 15 |
|  | Annealing | 58 | 15 |
|  | Elongation | 72 | 30 |
| 1 Cycle | Final elongation | 72 | 420 |
|  | Holding temp | 15 | ∞ |

Success of PCR was determined by electrophoresis using a 2 % (w/v) agarose gel stained with GelRed^®^ (Biotium, USA) and visualized under UV light. For SSR fragment size analysis, 2.0 μl – 3.0 μl of 2 different markers amplification products were co-loaded along with the internal size standard, GeneScan^™^ –500 LIZ^®^ (Applied Biosystems, USA) and Hi - Di^™^ Formamide (Applied Biosystems, USA). The DNA (PCR products) in this mixture was denatured for 3 minutes at 95 °C and chilled on ice for a few minutes.

The products were then separated by capillary electrophoresis using an ABI Prism^®^ 3730 Genetic analyzer (Applied Biosystems, USA) (Koumi et al., 2004). This provided automated and accurate estimates of allele sizes, which is better than using traditional gels because of the differences that can occur in migration between lanes in a gel (Life Technologies Corporation user guide, Carlsbad USA).

### Fragment Analysis

Fragment analysis was performed using Gene Mapper 4.0 (Applied Biosystems, USA) and allelic data for every marker was further analyzed by PowerMarker V3.25 (Liu and Muse, 2005) and DARwinV.6 (Dissimilarity Analysis and Representation for Windows^®^) software (Perrier and Jacquemound-Collet, 2006). Analysis of the molecular variance (AMOVA) was performed using Arlequin V.3.5.2.2. PowerMarker calculates a Table of summary statistics including inbreeding co-efficient, gene diversity and polymorphic information content (PIC), allele number; estimation of allelic and genotypic frequency, Hardy-Weinberg disequilibrium and linkage disequilibrium. PIC, which is a measure of diversity,was calculated using the formula:

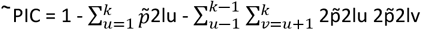

Where P_lu_ is the allele population frequency at the l^th^ locus and P_lv_ is the genotype population frequency at the l^th^ locus.

Dissimilarity was calculated by Darwin software using the formula:

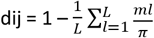

Where d_ij_ is the dissimilarity between units i and j, L is the number of loci, π is the ploidy and ml is the number of matching alleles for locus l.

DARwin was also used to display dendograms based on coconut evolutionary relationships using the dissimilarity matrix (Perrier and Jacquemound-Collet, 2006) as well as Principle Coordinate Analysis (PCOA) graphs

## RESULTS AND DISCUSSIONS

### Genotyping the coconut genotypes using capillary electrophoresis

Some common markers were used in the current study and that of Oyoo et al. (2016). The common markers used were CAC02, CAC03, CAC04, CAC06, CAC56, CAC72 and CN1C6, while CAC13, CAC20, CAC65, CN11E10, CN1G4, CN2A4 were specific to this study. Markers used by Oyoo et al. (2016) were CAC10, CAC21, CAC23, CAC71, CAC84, CN11E6 and CN1H2. In this study, 14 of the 30 markers failed to amplify DNA while three (3) were monomorphic. These were eliminated from the analysis. The markers that amplified well and produced bands across the 48 coconut genotypes are: CN1C6, CAC56, CAC03, CA08, CAC11, CAC20, CAC13, CAC39, CAC56, CAC10, CAC23, CAC72, CN11A10, CN11E10, CN11E6, and CN1G4 were amplified in all the 48 DNA samples as shown in Figure 1. Seven markers, CN1C6, CAC56, CAC03, CAC13, CAC20, CAC72 and CN1G4 produced visible bands though faint.

**Figure 1:**
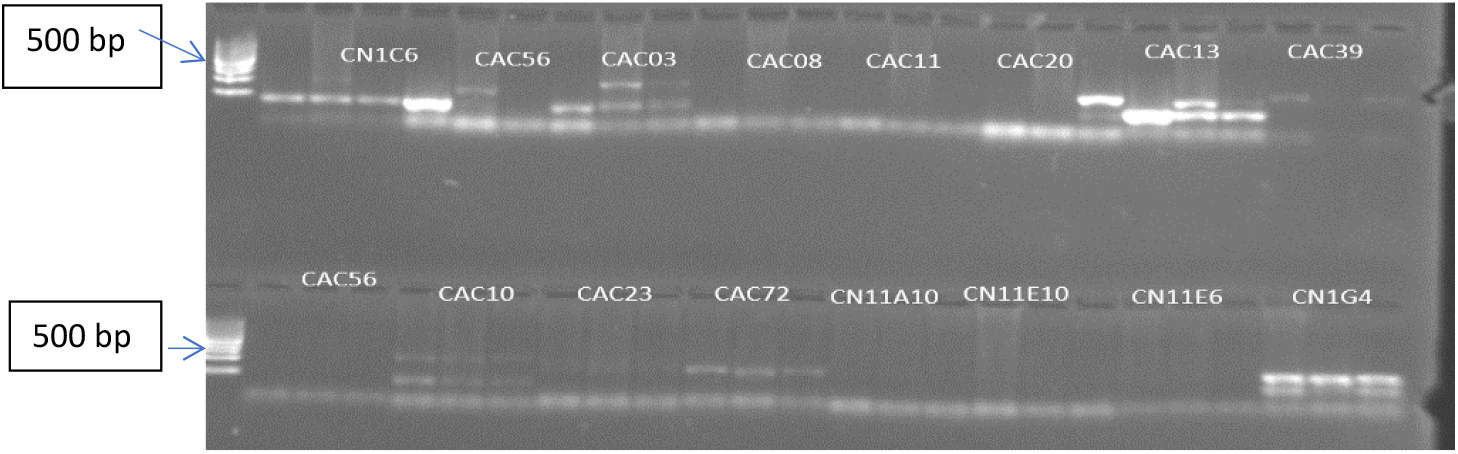
Agarose gel image (2 % w/v) of PCR products for SSRs markers that produced faint bands.

Figure 2 shows the bands from 7 polymorphic markers used to amplify three random coconut DNA samples using 2 % agarose gel electrophoresis using similar procedure as used by Oyoo et al., (2016). From this, it was not possible to tell by visual assessment alone whether the PCR products generated for each marker were similar or of different sizes. For example, for marker CAC20, the allele sizes indicated on the gel look the same (figure 2), but capillary electrophoresis determined that they were actually different band scores; 161 bp, 163 bp and 165 bp respectively. It should also be noted that CAC56, CAC03, CAC13, CAC20 and CAC72 displayed the co-dominance of SSRs markers, presenting two heterozygous alleles for some of the genotypes; this was not distinguishable in the previous work by Oyoo et al., 2016.

**Figure 2:**
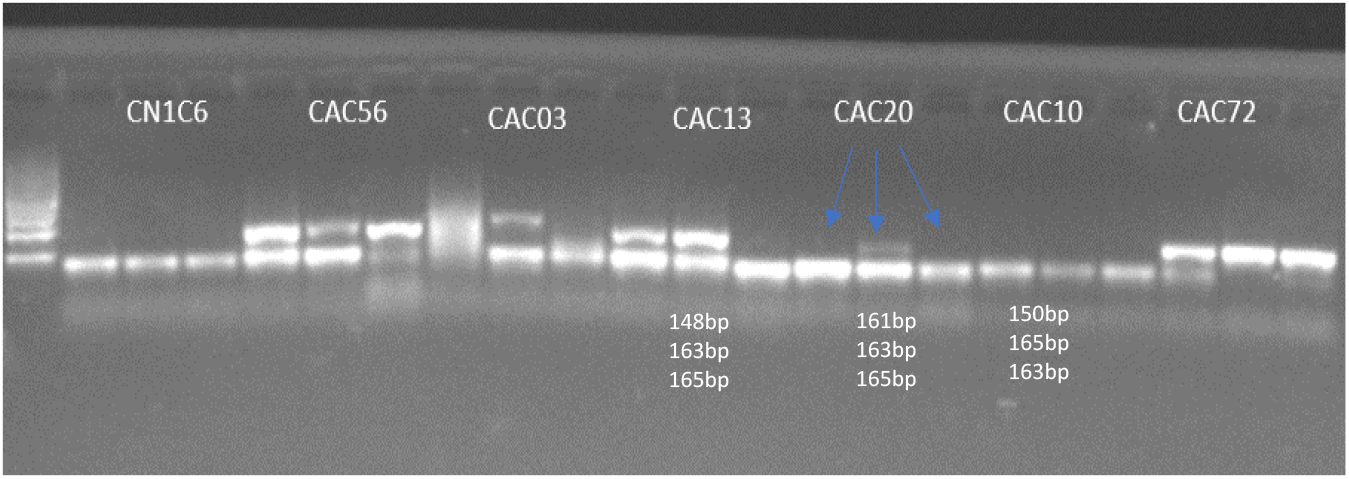
Agarose gel image (2.0 % w/v) of PCR products from 7 markers to confirm amplification of SSRs alleles prior to capillary electrophoresis.

Oyoo et al. (2016) assessed markers on the basis of presence or absence of the expected band. In contrast, in the current study, differences in allele sizes amplified for a single marker for the 48 coconut DNA samples were precisely assessed with capillary electrophoresis. For example, for CAC56, 6 possible genotypes were distinguished (Supplementary Table 1). Furthermore, none of the 13 polymorphic markers used in this study presented any absent alleles. Such absent PCR products should be considered carefully as they could have occurred as a result of failed PCR amplification or failure of primers to bind to the allele locus, and not necessarily due to an absent allele.

### Genomic diversity studies

The allelic data for the 13 SSRs markers was analyzed by PowerMarker^®^ version 3.25 and the summary statistics of allele frequencies, heterozygosity, polymorphic information content, gene diversity, allele number and number of genotypes discerned by each marker are presented in Table 3.

**Table 3:**
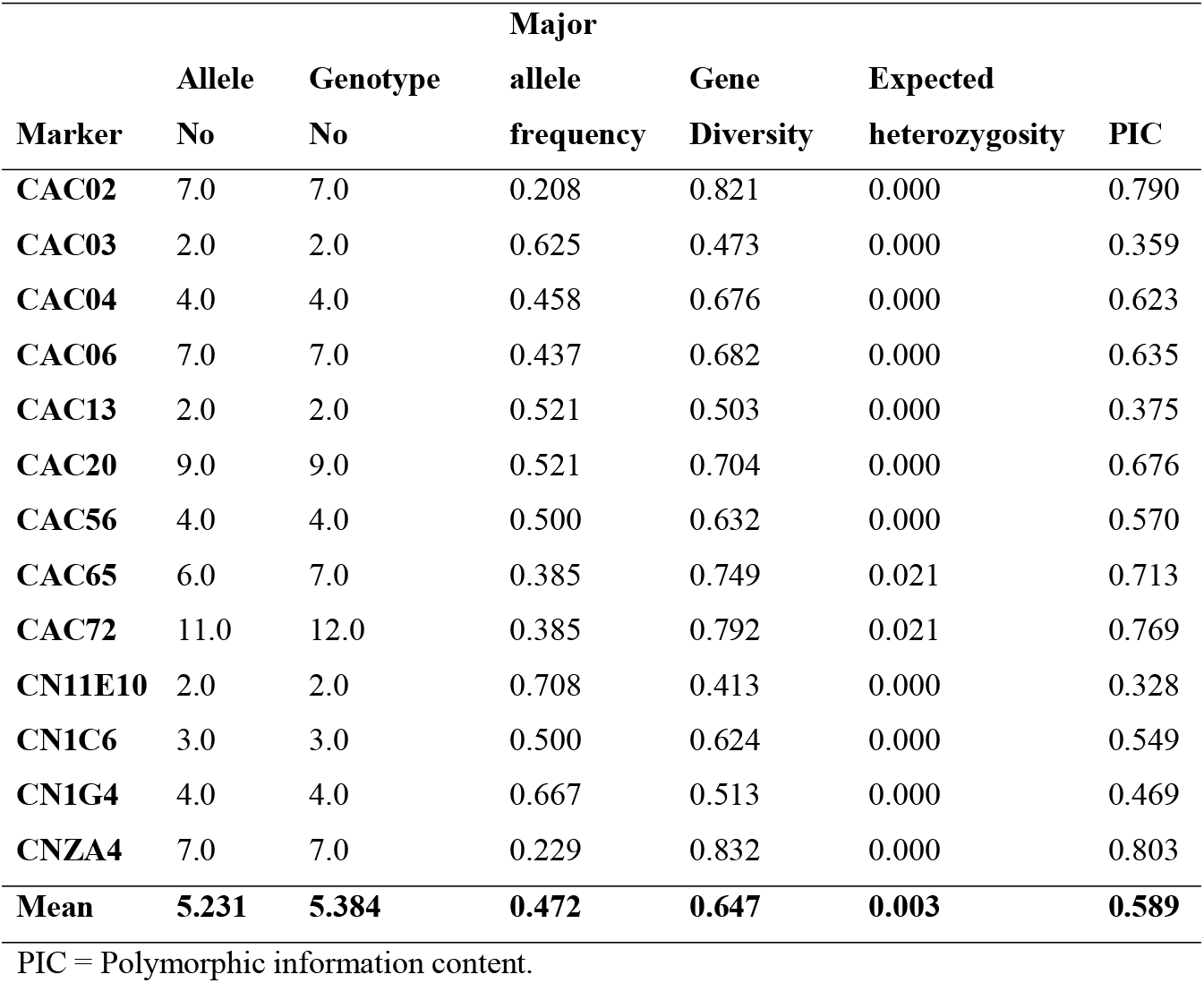
Summary statistics of allelic data analysis for 13 markers used to amplify the 48 coconut DNA samples in coastal Kenya.

### Polymorphism of the SSRs and Genetic Diversity of cape goose berry accessions

An average of 5 genotypes were detected by each marker in the study, marker CAC72 was the most sensitive because it differentiated the coconut population into twelve genotypes while markers CAC03, CAC13 and CN11E01 were the least sensitive differentiating only two genotypes. Overall, six SSR markers were able to differentiate the population into more than five genotypes and were considered to be sensitive (Table 3). The other six SSRs differentiated the population into either two, three or four genotypes only and were considered to be less sensitive.

In this study, a total of 68 observed alleles were detected, ranging from 2 for markers CN11E10, CAC13 and CAC03 to 11 for marker CAC72 with a mean of 5.231 alleles per marker. The numbers of alleles reported in the study are comparatively higher than those reported by Oyoo et al., 2016 using agarose gel electrophoresis. This finding informs that capillary electrophoresis has a higher resolution of detecting different alleles at a given locus than gel electrophoresis.

The highest major allele frequency was 0.708 (CN11E10) and the least was 0.208 (CAC02) with a mean 0.472 (maximum possible value is 1). This means that, using capillary electrophoresis at any given locus the mean chance of any of the alleles being detected is about 0.5 showing that this method has higher resolution for band size separation and no bias towards dominant alleles (Njung’e *et al*., 2013). This finding is in contrast with results of Oyoo *et al*., 2016, who reported a higher major allele values with a mean of 0.807 showing that gel electrophoresis has a lower resolution for band size separation and higher bias (80%) for detecting the dominant allele.

Polymorphic information content (PIC), a measure of how well the marker distinguished the samples tested, ranged from 0.33 for marker CAC03 to 0.80 for marker CNA4 with a mean of 0.59.This was in contrast to Oyoo *et al*., 2016 who reported who reported the highest PIC of 0.364 with a mean of 0.235. Six primers CNZA4 (0.8026), CAC02 (0.7904), CAC72 (0.7697), CAC04 (0.6228), CAC06 (0.6346), CAC20 (O.6763) and CAC65 (0.7134) showed higher PIC values. These primers should therefore be given priority in genotyping coconut because they have higher segregation capacity. The PIC values showed in this study were significantly higher compared to Oyoo et al., 2016. This shows that capillary electrophoresis significantly increases the efficacy of SSR markers and therefore increases their resolution for detecting diversity in coconut genotypes. This suggests that these markers are suitable for detecting the genetic diversity of coconut accessions from the Coast of Kenya.

Expected Heterozygosity for the selected markers was generally low with a mean of 0.003 (minimum possible value is 0) indicating that the materials tested were genetically pure, i.e. the loci were stable and not prone to high outcrossing frequencies. These markers were good for diversity studies especially in the tall coconut varieties, due to their outcrossing nature (Oyoo et al., 2016).

The study reaffirms that SSRs are powerful tools in genetic diversity studies and their usefulness could be enhanced if capillary electrophoresis was used than the standard procedure. In DNA markers diversity studies, allelic data analysis is the key towards attaining conclusive and reliable results (Vemireddy, 2007). The choice of allele sizing and separation platforms determines the accuracy in allele sizing, reproducibility and interpretation of the data obtained (Wang et al., 2009). A comparative analysis across 8 microsatellite loci in 12 rice varieties (Vemireddy, 2007) demonstrated that capillary electrophoresis is the most accurate and preferred method for allele size estimation, with errors of less than 0.73 bp compared to slab gels such as polyacrylamide, which produce error rates of up to 1.59 bp and agarose gels, which give error rates of up to 8.03 bp. Capillary electrophoresis has also been proven to have greater reproducibility (3 bp). Oyoo et al. (2016) who used 2 % (w/v) agarose gels could not separate accurately alleles with size differences of as little as 2 bp where the average allele length is 150 to 500 bp long. Such alleles will migrate the same distance in an agarose gel and will therefore be assumed to be of the same size leading to limited interpretation. SSRs results should not be scored as present or absent as they are co-dominant markers and missing alleles can be as a result of low quality DNA or due to a marker that could not bind specifically to the allele locus. In this study, capillary electrophoresis was used to separate alleles and hence added value to the previous study by Oyoo et al., (2016).

### Population structure of Coconuts (*Cocos nucifera* L) along the coast of Kenya

#### Genetic distances of coconut genotypes among counties

The results of partitioning of genetic variance within and among coconut populations sampled presented in Tables 4 and 5. The AMOVA was performed using Arlequin software. Samples obtained from four Counties; were considered as constituting 4 different populations.

**Table 4:** AMOVA results of four coconut populations from the coastal lowlands of Kenya.

| Source of variation | Degrees of freedom | Sum of squares | Variation components | Percentage of variation |
| --- | --- | --- | --- | --- |
| Among populations | 3 | 21.265 | -0.06779 | -1.59 (p = 0.001) |
| Among individuals within populations | 44 | 380.402 | 4.31233 | 101.10 (p = 0.999) |
| Within individuals | 48 | 1.000 | 0.02083 | 0.49 (p = 0.001) |
| Total | 95 | 402.667 | 4.26538 |  |
| Significance level (0.05) |  |  |  |  |

**Table 5:**
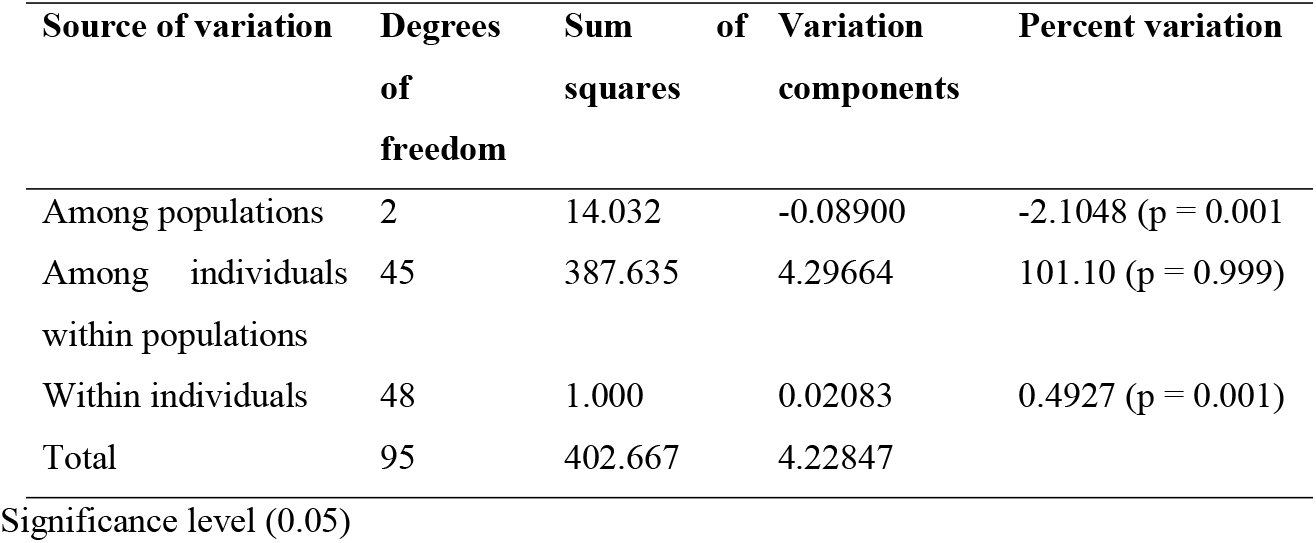
The analysis of molecular variance (AMOVA) of the coconut germplasm in the coastal lowlands of Kenya

Results of the study showed a significant negative variation −1.59% (p = 0.001), in coconut genotypes in different counties indicating lack of population structuring according to county of collection. While the population molecular variance value reported in this study was lower than that reported by Oyoo et al, 2016 (2%) who used the same markers and gel electrophoresis detection platform, both studies reported lack of population differentiation.

The lack of population differentiation may be attributed to a number of factors; the nature of markers used may lack enough resolution to group the genotypes according to the county of collection (Excoffier et al., 2005). There are chances that similar genotypes were sampled between counties or there is exchange of germplasm by farmers between Counties. The negative variance can also be attributed to the highly outcrossing nature of coconut (Gunn et al., 2011) resulting in exchange of genes between populations/counties and reducing interpopulation diversity compared to within same population (Excoffier et al., 2005).

Overall, no geographical divergence was observed in coconuts evaluated in the study across the selected counties. This result shows that coconut populations in the select counties at the Kenyan coast are similar.

#### Genetic variations among coconut genotypes within counties

The percentile variation among individuals within counties was not statistically significant 101.10 (p = 0.999) indicating that the differences observed in the coconuts within counties could be due to a shared pedigree within the populations. The percentage variation within individuals in the same county was significant 0.4927 (p = 0.001) indicating existence of different distinct genotypes in the counties. The molecular individual variance was statistically significantly higher than population variance in the study.

Overall, the coconut genotypes within the select Kenyan coastal counties are genetically diverse. It can therefore be inferred that diversity of the coconut in the select counties is as a result of genetics and not geography.

#### Factorial analysis

Factorial analysis was performed to evaluate the genetic relationship of the genotypes within and between the counties. The scatter plot was derived from the dissimilarity matrix calculated using raw binary data using Darwin 6.0.21 software. Factorial analysis failed to group the genotypes into any discernible pattern either based on county of collection or genetic makeup (Figure 3). This observation further confirms that the technology applied was limited in its resolution of segregating the coconut genotypes.

**Figure 3:**
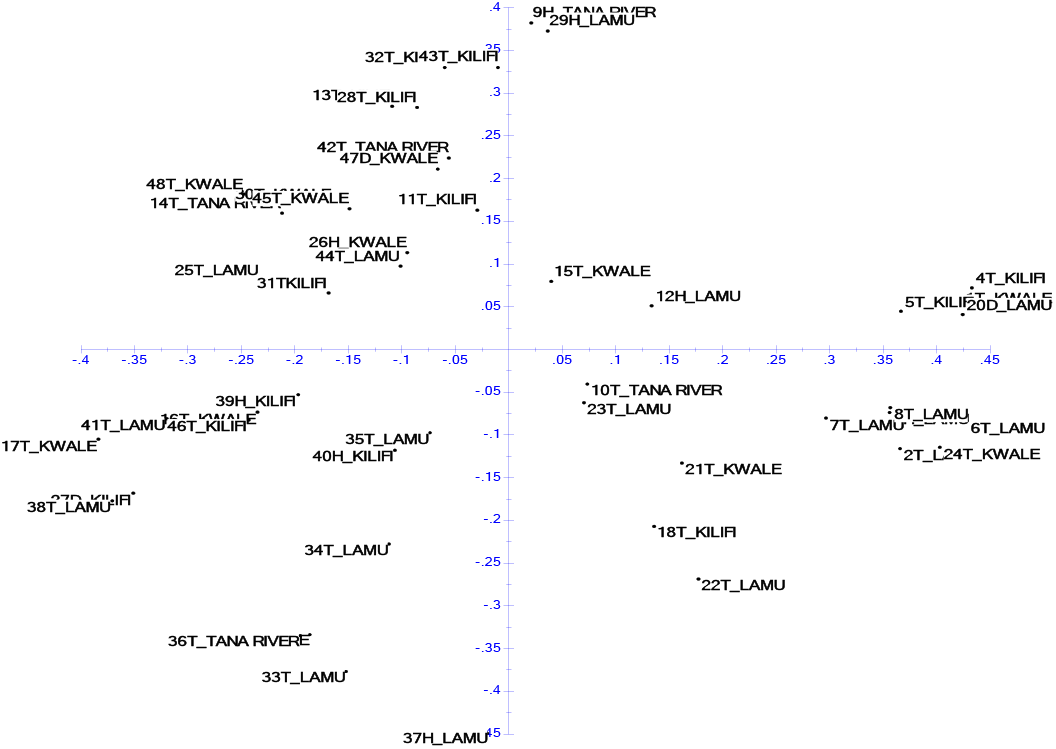
Factorial analysis of the coconut genotypes from select Kenyan coast counties

#### Phylogenetic analysis

Relationship of the genotypes was presented in a Neighbor Joining dendrogram derived in Darwin 6.0 based on Sokal – Michener modalities after one thousand bootstraps (Figure 4). Phylogenetic analysis grouped the genotypes into three main clusters and eleven sub clusters randomly without any specific pattern. Analysis of markers did not segregate the genotypes based on either genetic makeup or county of collection. This observation further highlights the inefficiency of the markers used in detecting diversity within the coconut genotypes in the four select counties of the Kenyan coast. The lack of genotype structuring may be due to genetic closeness of the coconut genotypes and or low resolution of the markers used in the study. This finding is similar to results of Oyoo et al., 2016 who also failed to achieve segregation of the genotypes using the same markers.

**Figure 4:**
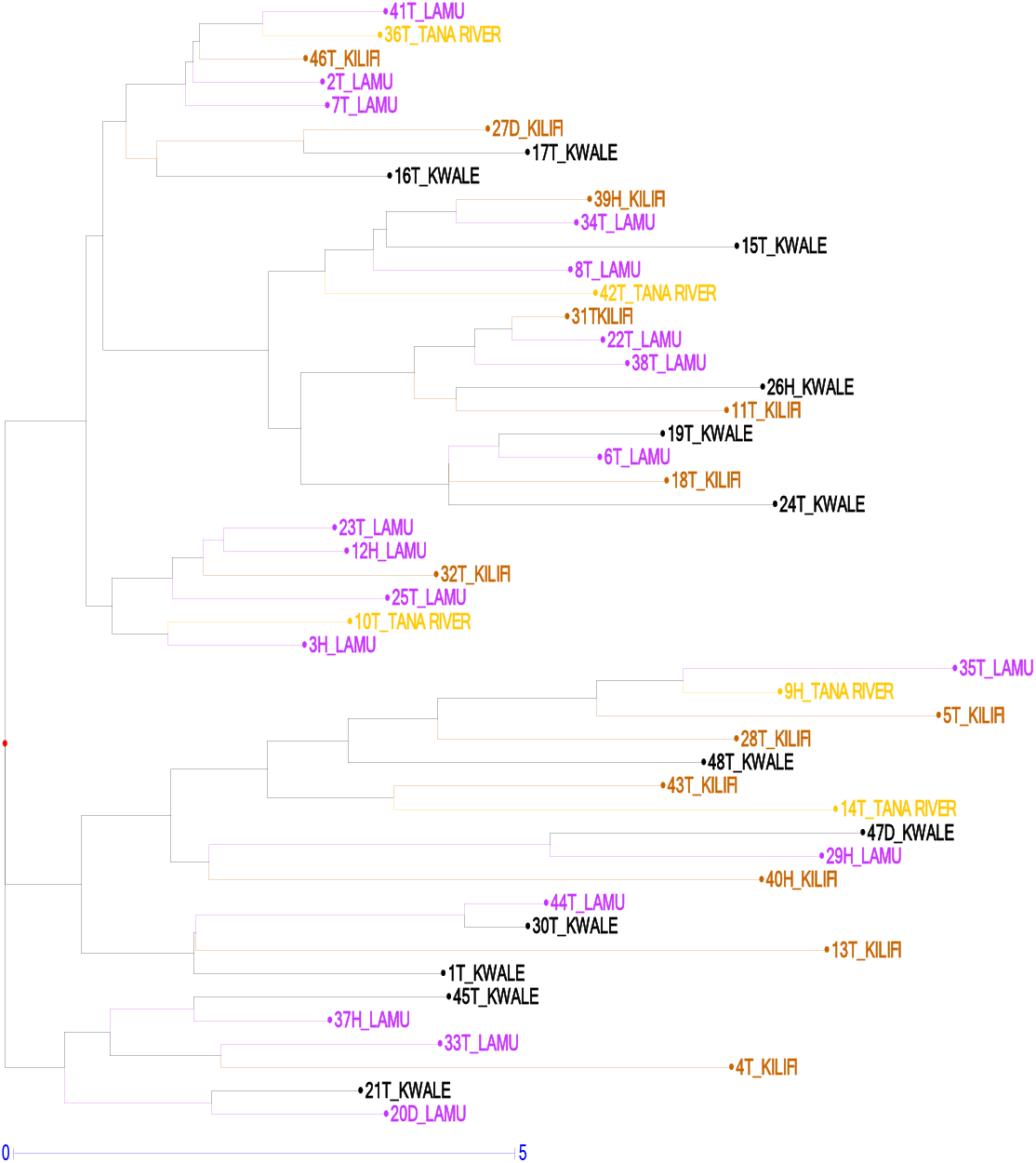
Neighbour-joining DENDOGRAM showing patterns of clustering of the coconut genotypes in the four counties of coastal Kenya. *Genotypes are represented by numerical identities followed by a letter designating the coconut variety; T=Tall, D=Dwarf, H=Hybrid*.

The lack of population partitioning is common in free mating populations where alleles are freely shared within and populations. This postulation is further confirmed by the low expected heterozygosity values reported in the study (Table 3). The lack of population differentiation can also be attributed to high cultivation and domestication coconut leading farmers transferring germplasm within and across the counties.

## CONCLUSION AND RECOMMENDATIONS

This study demonstrated that capillary electrophoresis is a superior method for DNA length polymorphisms detection among the studied germplasm as compared to agarose gel electrophoresis. This new information was used to enhance the existing description of the diversity of the genotypes evaluated and guide establishment of a coconut genebank for *ex-situ* conservation.

The method still established the narrow genetic base of Kenyan coconut germplasm suggesting a need to expand coconut genetic base in Kenya, through introductions and controlled crosses of elite parents

DNA based genetic characterization will also help avoid duplications in conservation blocks and breeding programs, maximising the use of genetic diversity between coconut populations to breed superior varieties and to identify coconut populations with narrow genetic bases and adoption of appropriate conservation strategies (Martial et al., 2013; Rao and Tobby, 2004).

## ACKNOWLEDGEMENT

National Commission for Science, Technology and Innovation (NACOSTI) and Kenya Coastal Development Project (KCDP) for financial support and the International Crops Research Institute for the Semi-Arid Tropics (ICRISAT) for technical facilities and support.

---

**Supplementary Table 1:**
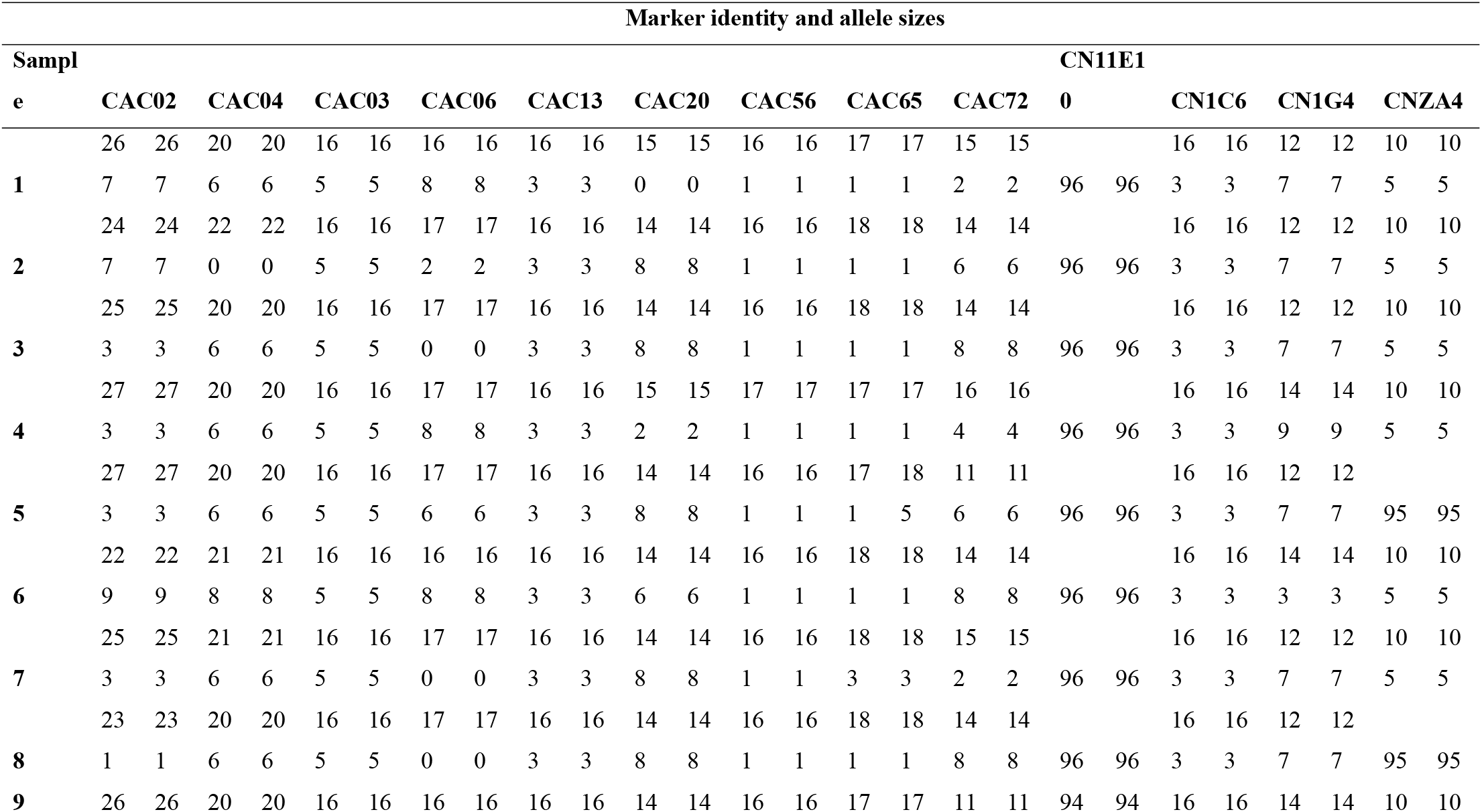
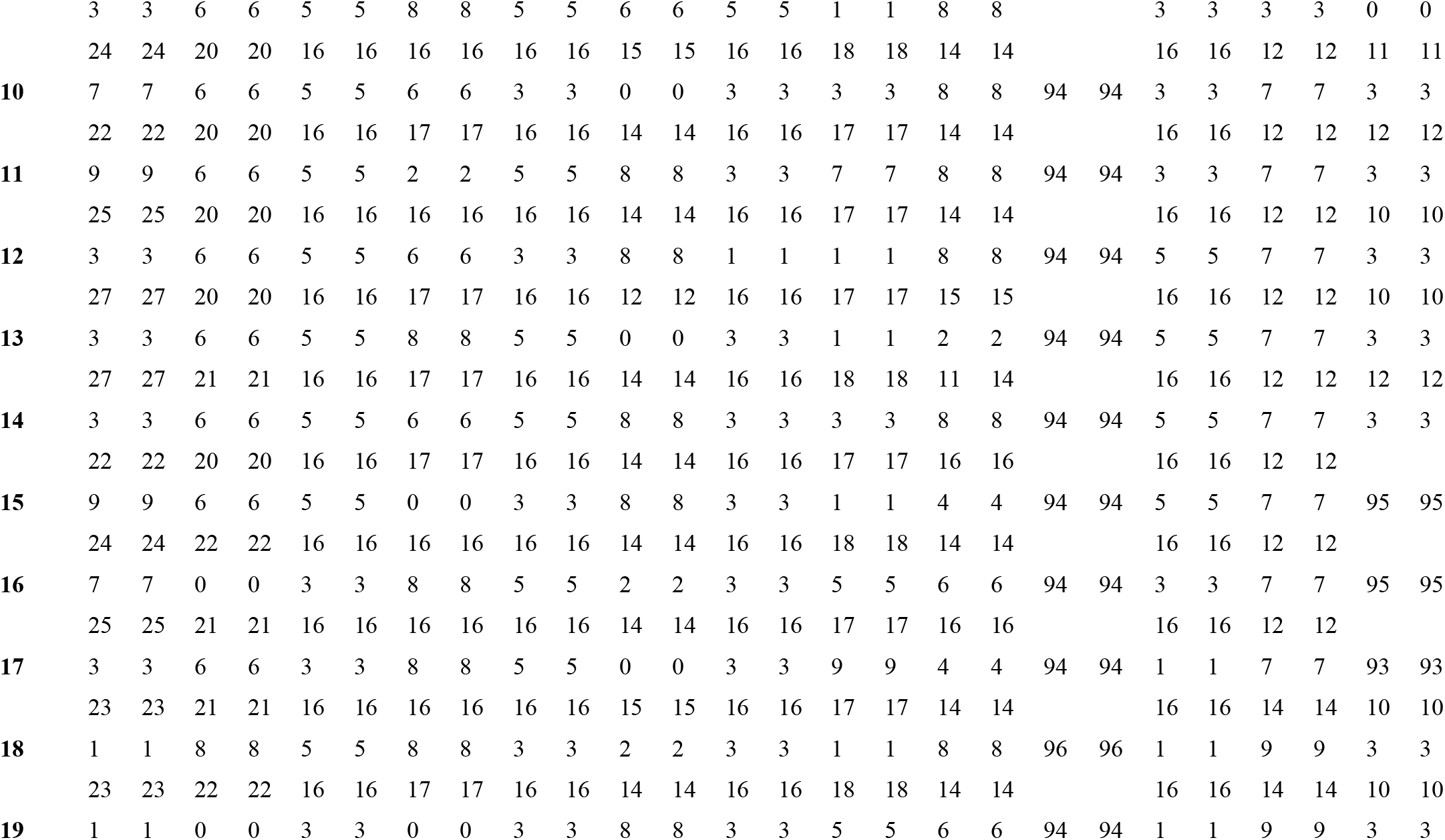
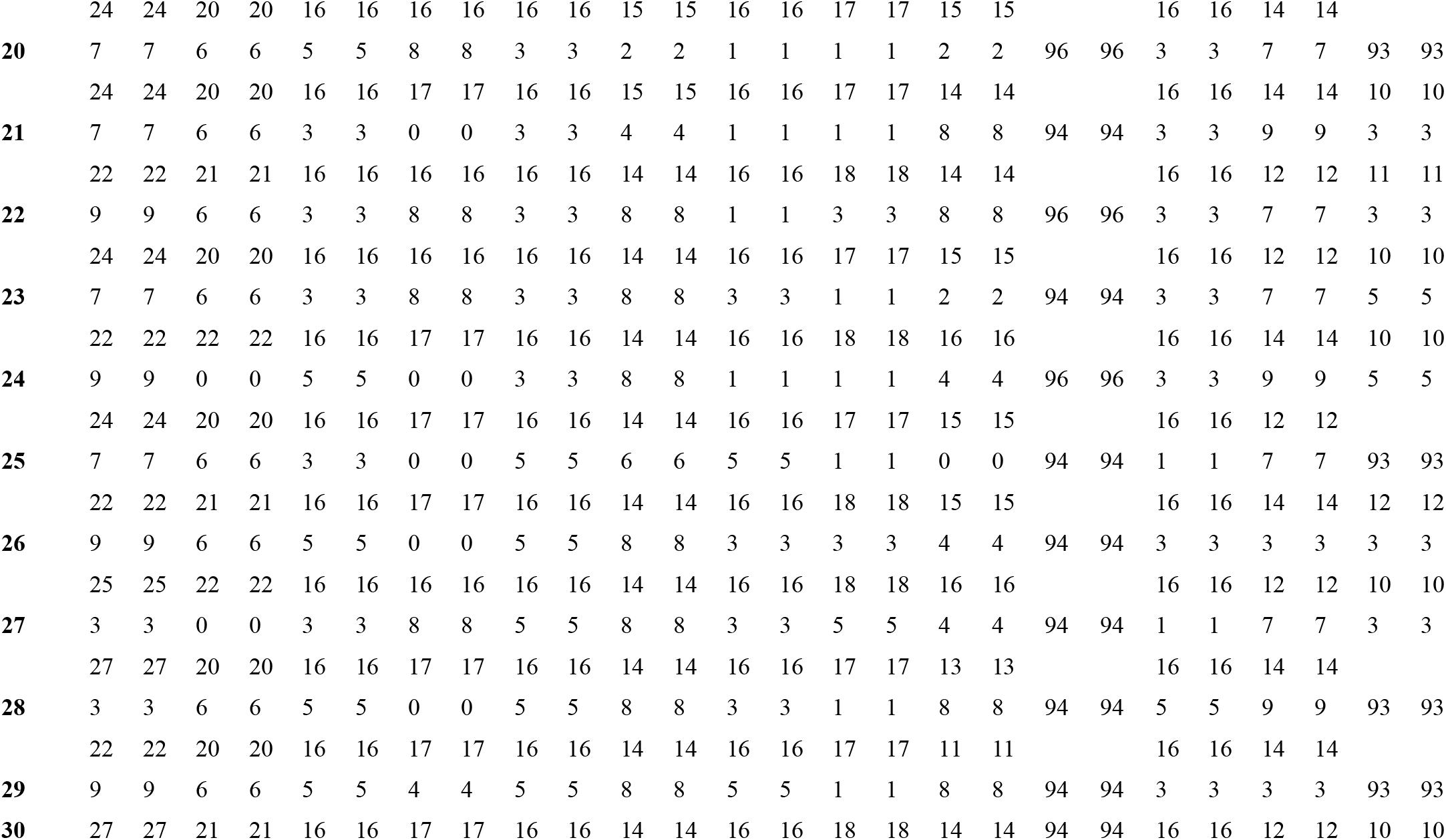
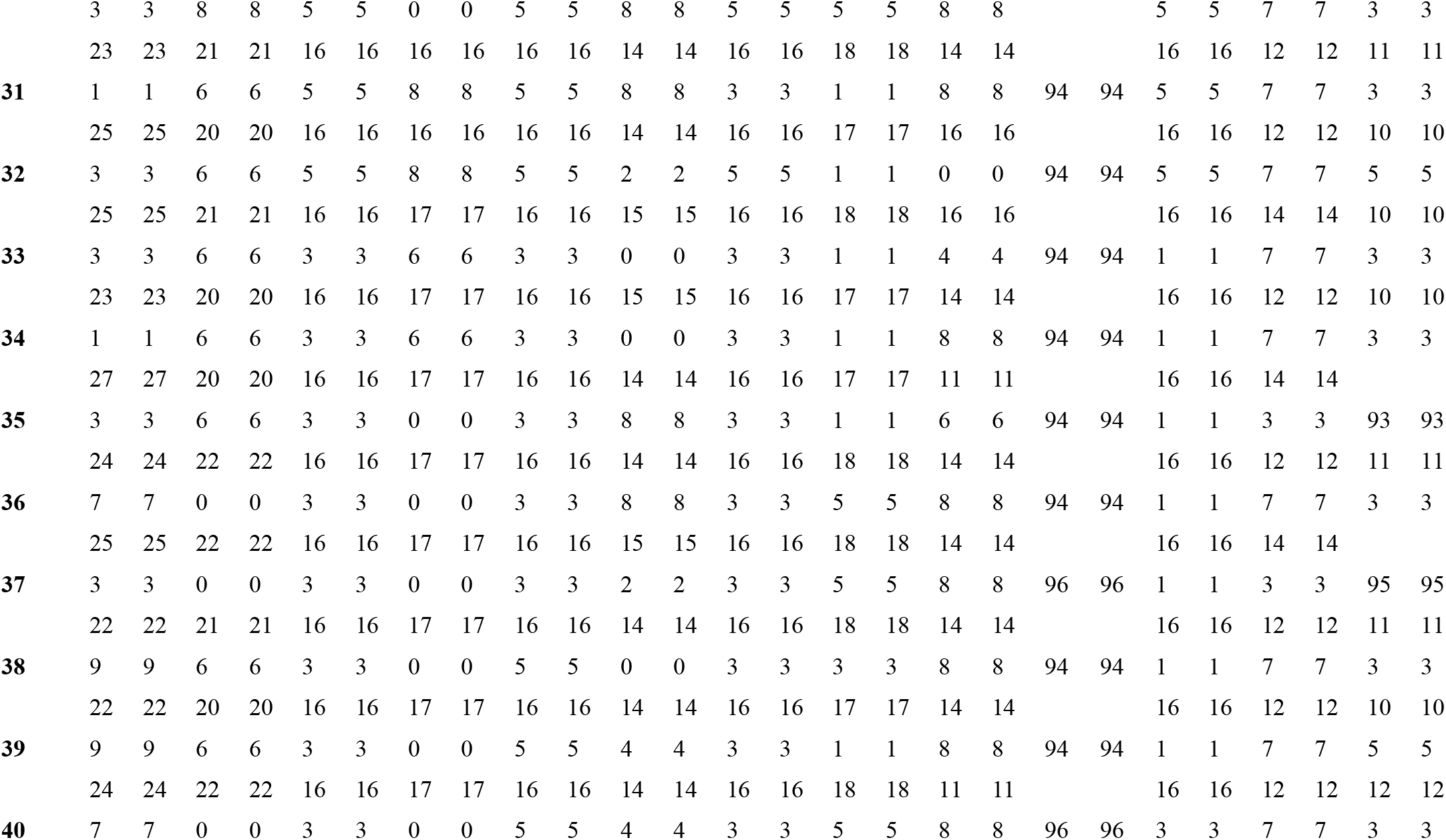
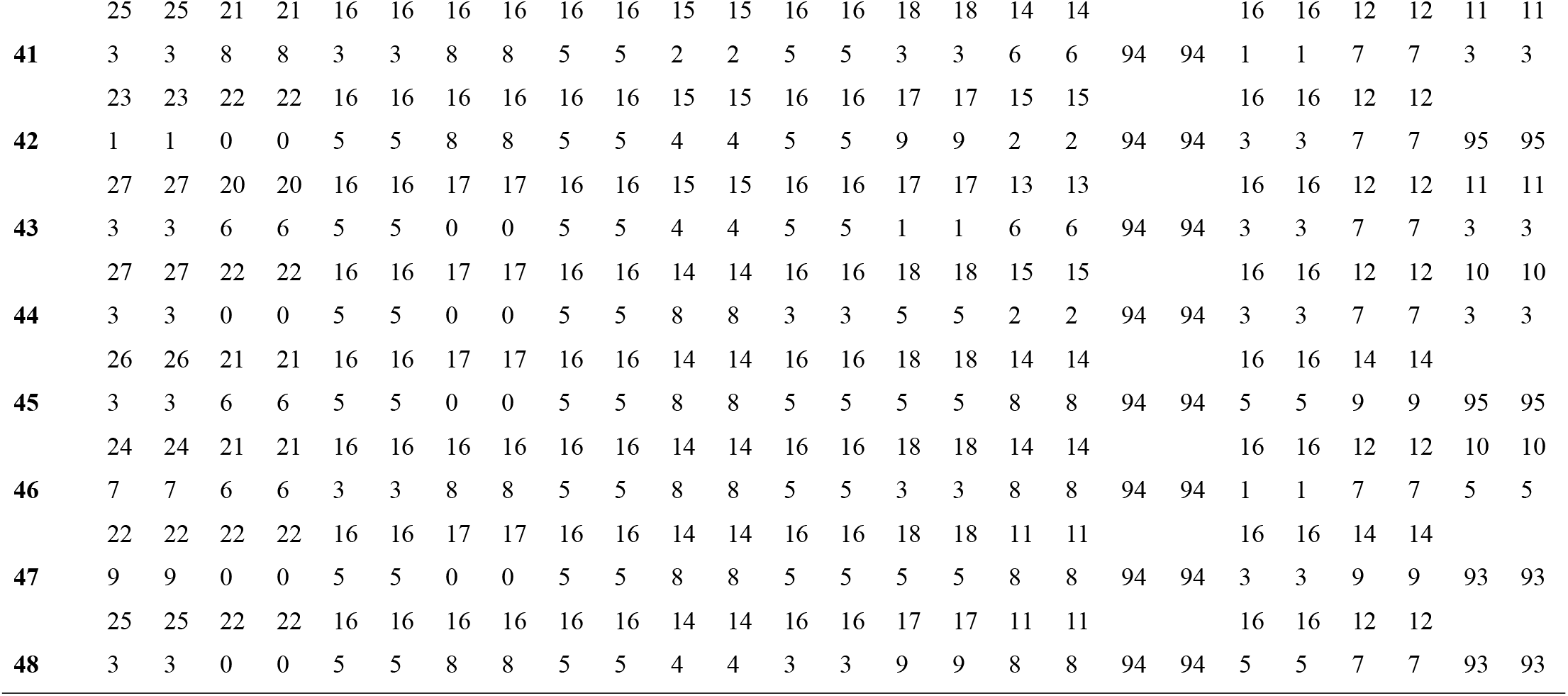
Genotyping data of 48 coconut genotypes and 13 polymorphic SSRs markers.

**Supplementary Table 2:**
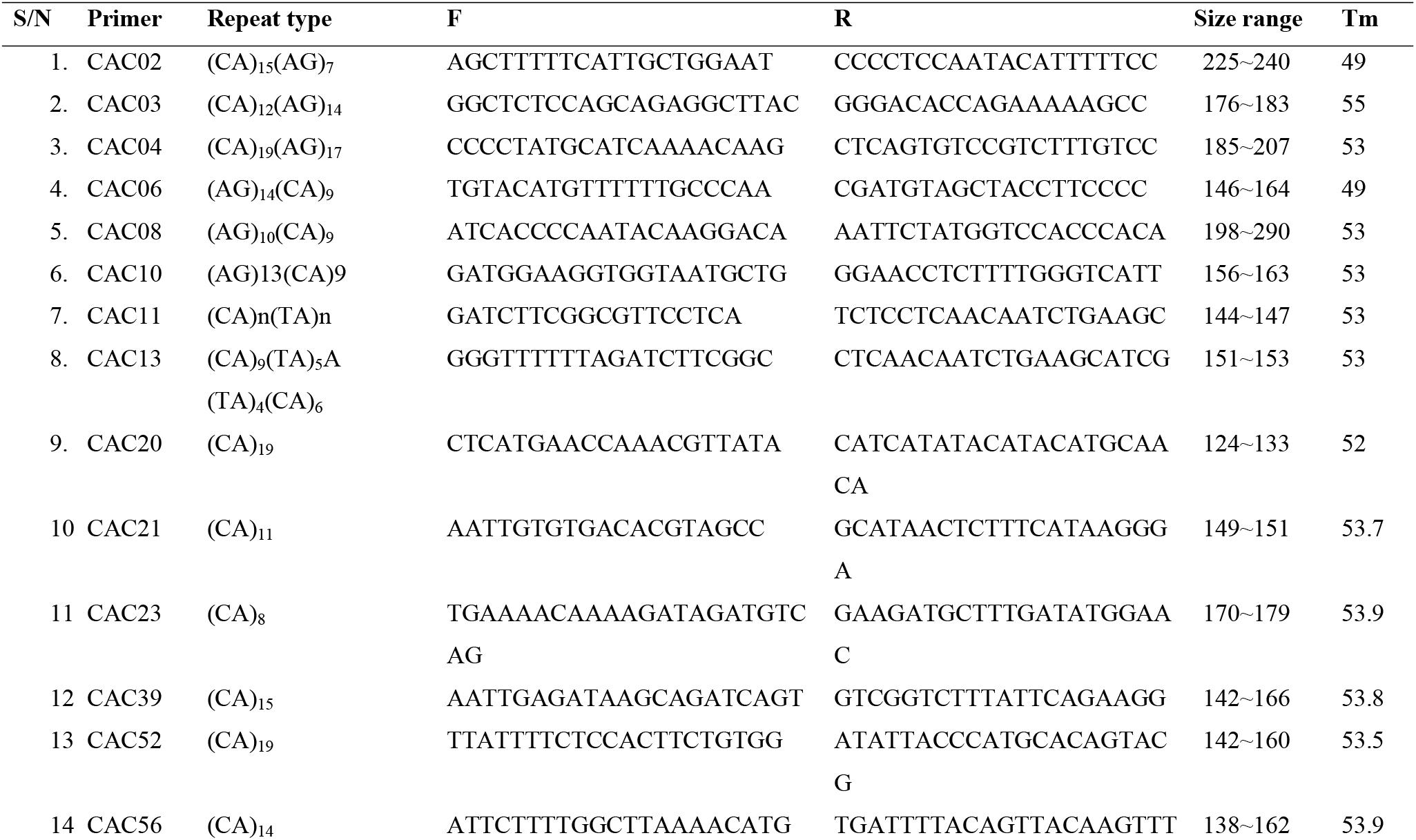
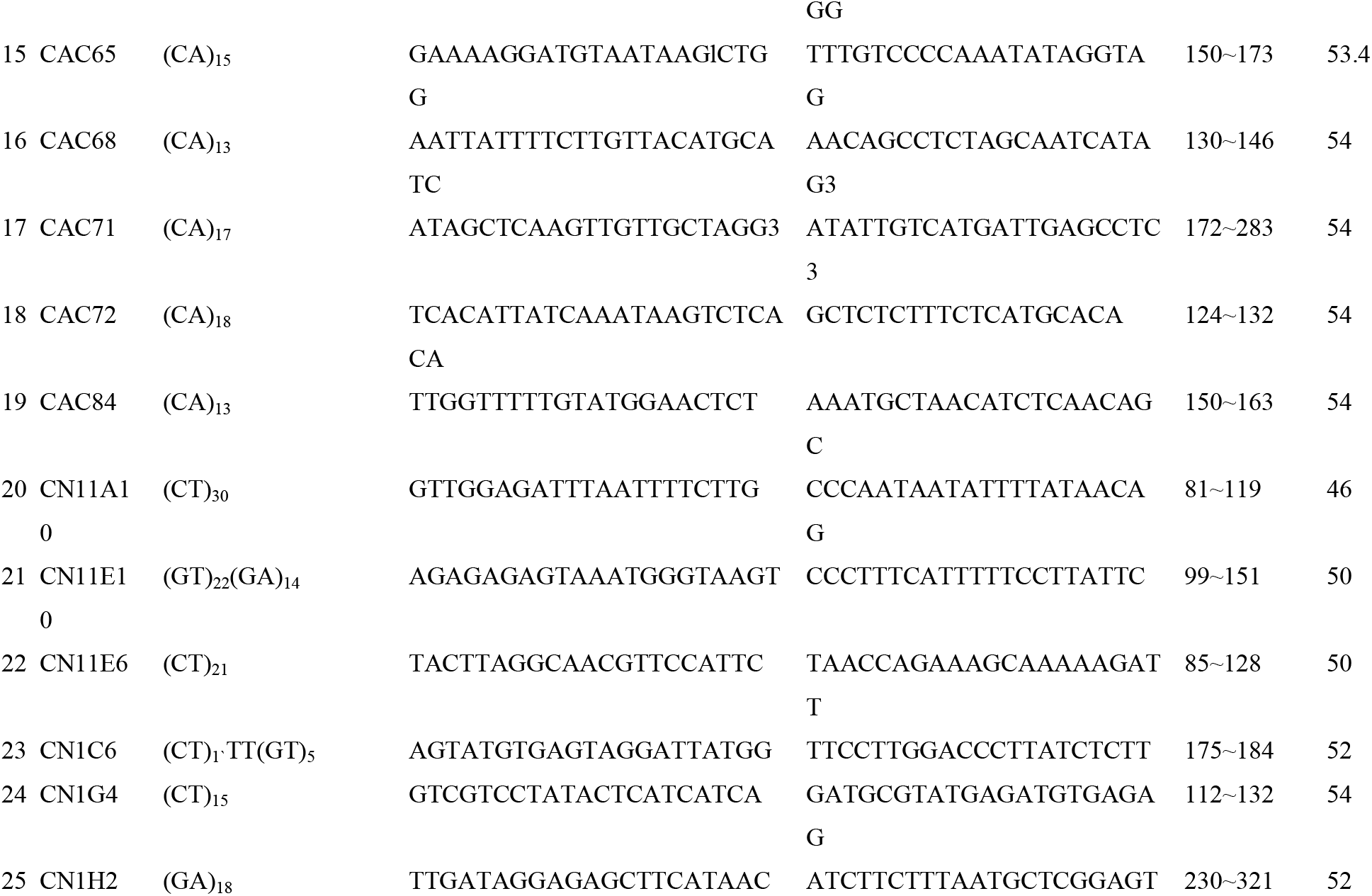
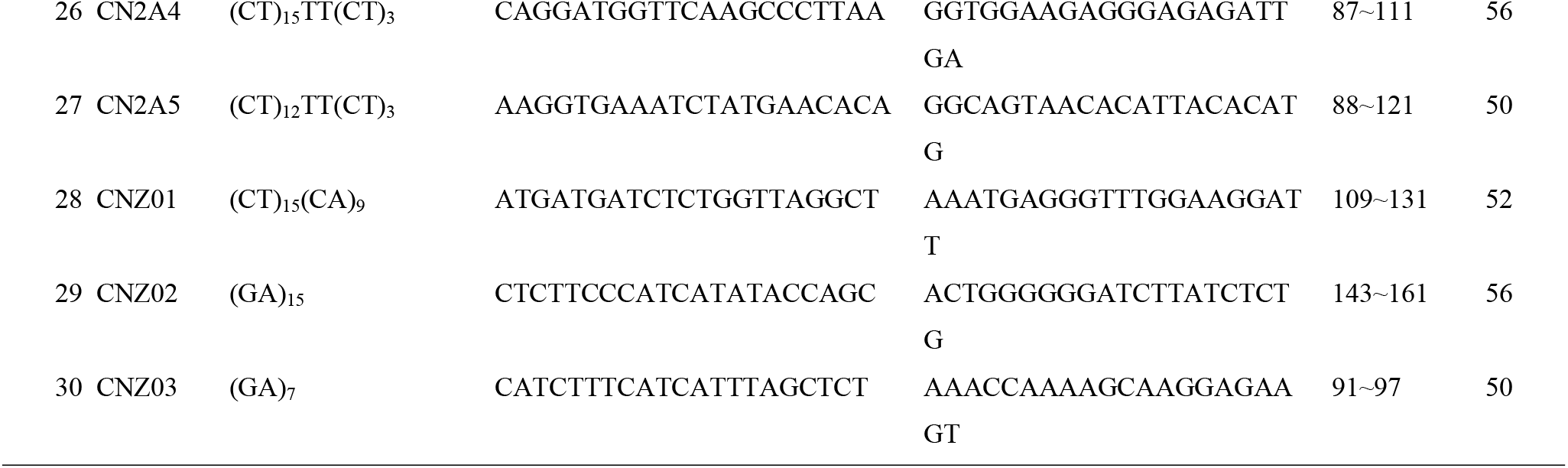
List of 30 SSR primers used in the study.

